# A review of co-occurrence and hybridization as neglected factors in studies of Syrian Woodpecker *Dendrocopos syriacus* and Great Spotted Woodpecker *Dendrocopos major*

**DOI:** 10.1101/2024.07.25.604904

**Authors:** Antonii Bakai, Gerard Gorman, Łukasz Kajtoch

## Abstract

Some species of woodpecker (Picidae), such as in the genus *Dendrocopos*, are known to occasionally hybridize. The distribution, biology and ecology of the Syrian Woodpecker (*D. syriacus*) and the Great Spotted Woodpecker (*D. major*) are fairly well-known (less so in the case of Syrian), but these closely related species are seldom treated together in studies. This review summarizes the published data on these species in order to evaluate the omissions and inaccuracies in research and surveys on their sympatric populations. As research that deals with both species together is scant, the need to examine interactions, both antagonistic and hybridization, is advisable in order to properly understanding their ecology, ethology, breeding biology and demography.

## 1. Introduction

In recent decades, woodpeckers have started to attract an increasing amount of attention from researchers, as they are considered to be a group of great significance in avian conservation biology owing to their role as keystone species, “ecosystem engineers” and indicators of habitat quality (Jones, Lawton & Schachak 1994; Martin 2015; Menon & Shahabuddin 2021). This group of birds are often important keystones in urban environments, where natural cavities in trees can be limited (Catalina-Allueva & Martín 2021). Hence, our understanding of their biology is crucial for proper environmental management and conservation plans and projects. Among the European woodpecker species which occur in “secondary habitats”, in urban, suburban and rural environments, particular interest has arisen around two closely related species – Great Spotted Woodpecker (GW) and Syrian Woodpecker (SW), as they are often sympatric in occurrence and are known to compete but at the same time will form mixed pairs and produce viable and fertile offspring (Gorman 1999; Dudzik & Polakowski 2011; Kajtoch & Kusal 2022). These two species are members of a taxonomic complex that includes Sind (*D. assimilis*), White-winged (*D. leucopterus*), Himalayan (*D. himalayensis*), and Darjeeling (*D. darjellensis*) woodpeckers, which occur outside Europe in central and/or southern Asia. Genetically, these woodpeckers are also related to the White-backed Woodpecker (*D. leucotos*), which occurs across the Palearctic, and to the Okinawa Woodpecker (*D. noguchii*), which only occurs on the island of Okinawa, Japan. The phylogenetic relationships of these species are as follows: ((leucotos, noguchii),((major, leucopterus),((syriacus, assimilis),(himalayensis, darjellensis)))) (Fuchs & Pons 2015). Hybridization has been reported for most of these species when they occur in sympatric or parapatric populations. GW and SW have some distinct biological and ecological characteristics (habitat preferences, nesting sites), but in other features (morphology, breeding) they are also remarkably similar (see Table 1 and Fig. 1).

**Fig. 1.**
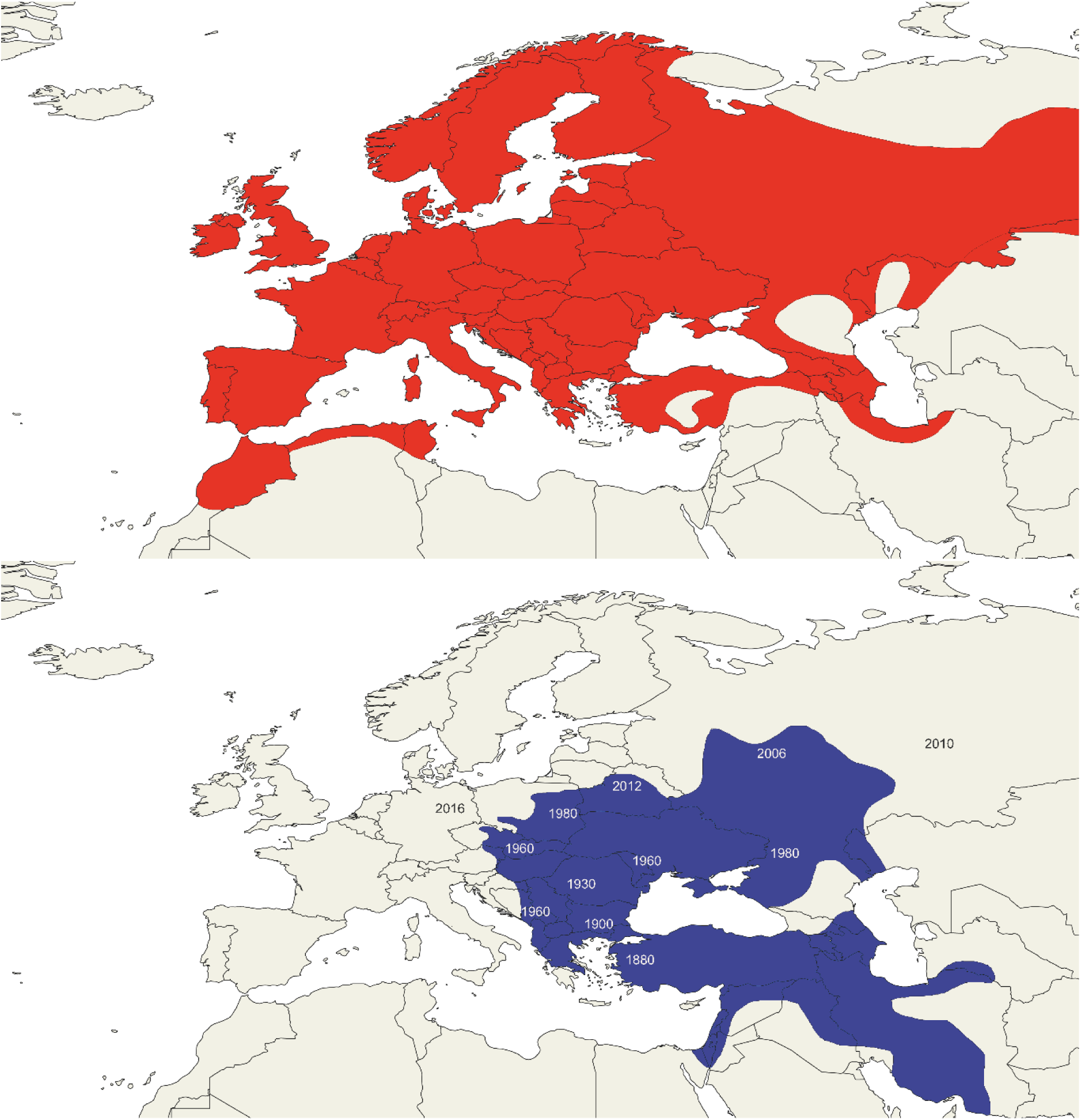
Ranges of Great Spotted (above) and Syrian (below) Woodpeckers in the Western Palearctic. Years on the Syrian Woodpecker map indicate the approximate dates of arrival in the given areas.

**Table 1.**
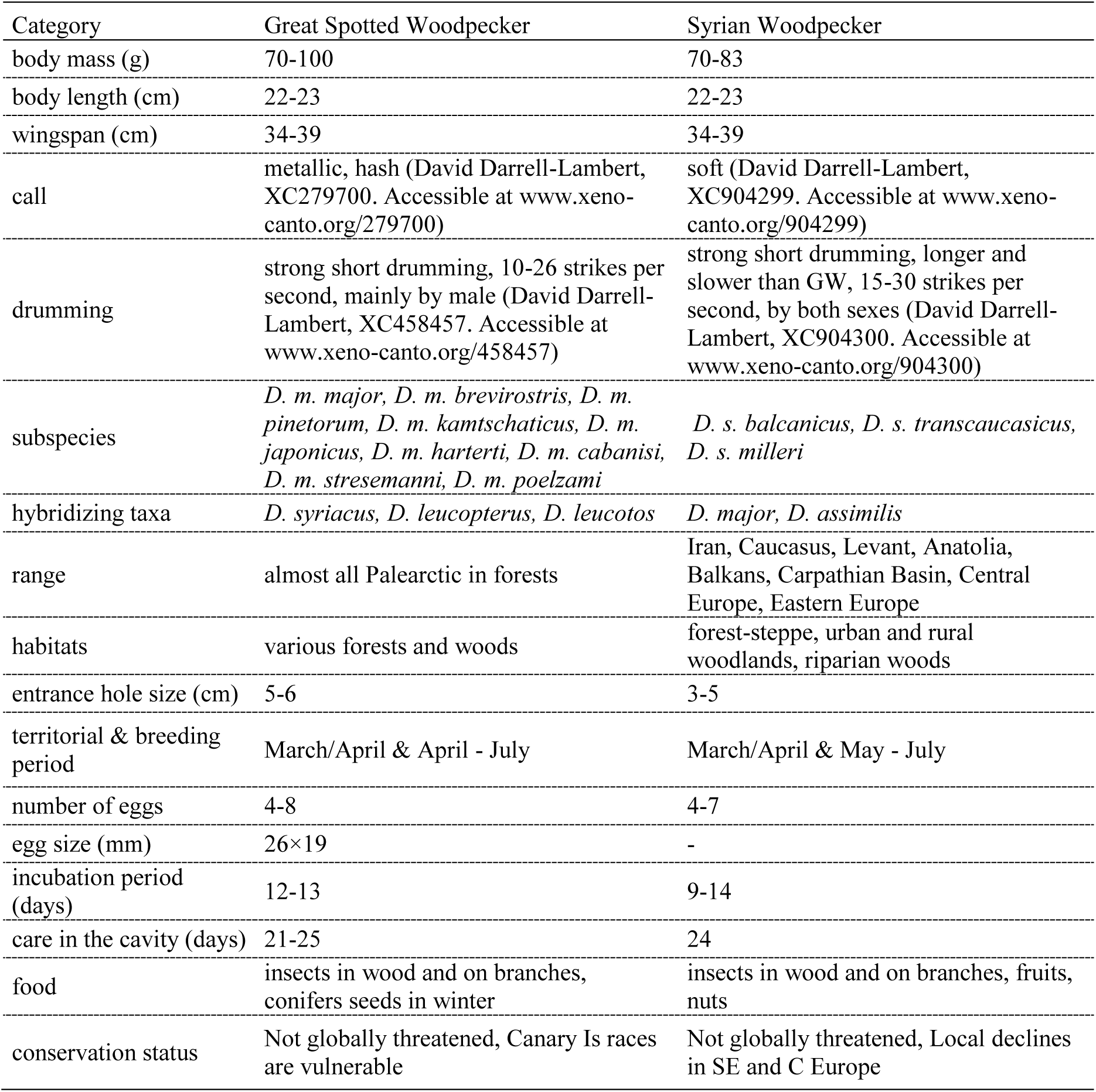
Characteristics of great-spotted and Syrian woodpeckers.

| Category | Great Spotted Woodpecker | Syrian Woodpecker |
| --- | --- | --- |
| body mass (g) | 70-100 | 70-83 |
| body length (cm) | 22-23 | 22-23 |
| wingspan (cm) | 34-39 | 34-39 |
| call | metallic, hash (David Darrell-Lambert, XC279700. Accessible at www.xeno-canto.org/279700) | soft (David Darrell-Lambert, XC904299. Accessible at www.xeno-canto.org/904299) |
| drumming | strong short drumming, 10-26 strikes per second, mainly by male (David Darrell-Lambert, XC458457. Accessible at www.xeno-canto.org/458457) | strong short drumming, longer and slower than GW, 15-30 strikes per second, by both sexes (David Darrell-Lambert, XC904300. Accessible at www.xeno-canto.org/904300) |
| subspecies | *D. m. major*, *D. m. brevirostris*, *D. m. pinetorum*, *D. m. kamtschaticus*, *D. m. japonicus*, *D. m. harterti*, *D. m. cabanisi*, *D. m. stresemanni*, *D. m. poelzami* | *D. s. balcanicus*, *D. s. transcaucasicus*, *D. s. milleri* |
| hybridizing taxa | *D. syriacus*, *D. leucopterus*, *D. leucotos* | *D. major*, *D. assimilis* |
| range | almost all Palearctic in forests | Iran, Caucasus, Levant, Anatolia, Balkans, Carpathian Basin, Central Europe, Eastern Europe |
| habitats | various forests and woods | forest-steppe, urban and rural woodlands, riparian woods |
| entrance hole size (cm) | 5-6 | 3-5 |
| territorial & breeding period | March/April & April - July | March/April & May - July |
| number of eggs | 4-8 | 4-7 |
| egg size (mm) | 26×19 | - |
| incubation period (days) | 12-13 | 9-14 |
| care in the cavity (days) | 21-25 | 24 |
| food | insects in wood and on branches, conifers seeds in winter | insects in wood and on branches, fruits, nuts |
| conservation status | Not globally threatened, Canary Is races are vulnerable | Not globally threatened, Local declines in SE and C Europe |

While GW has been widely studied, its congener SW “*can be considered the least known species among the European woodpecker species* (…)” and “*none of* [publications] *reported on any of the vital rates* (…)” (Pasinelli 2006).

Across much of their respective geographical ranges, GW and SW do not coexist. GW inhabits a wide range of wooded habitats across the entire Palearctic region, whilst the range and habitat use of SW is more restricted. However, where the geographical ranges of the two overlap, they sometimes occupy different habitats and altitudes. For example, in some areas, SW is often found in parks, gardens and natural open woodlands in lowlands, while in contrast GW inhabits denser closed forests, often in uplands (Gorman 2004) ) (Fig. 1). However, at the end of the 19^th^ century, the SW began a rapid range expansion in south-east Europe (Glutz von Blotzheim & Bauer 1980; Michałczuk 2014), occupying the Balkans and then the Carpathian Basin, before continuing into parts of central and eastern Europe. The species began the expansion from its original, native range in Iran, Iraq, the Levant and Anatolia into Europe at the end of the 19th century (Glutz von Blotzheim & Bauer 1980; Michalczuk 2014). It first colonised new areas in the Balkans before gradually moving westwards. It reached Slovenia in 1974 (Bacani 1998) and moved rapidly northwards onto the Great Hungarian Plain and the Wallachian Plain. From this areas it continued its colonisation northwards through Slovakia to Poland and through Austria to South Moravia in Czechia. In the east, it spread through Ukraine and eastern Russia before reaching the River Volga (Suravenkov 2022, Moskvichev 2020). An isolated population exists in Kazakhstan (Keller et al, 2020). The current northernmost edge of its distribution is on Baltic Coast in Poland and areas to the north of Moscow in Russia (Yankevich & Koshelev 2022). The origins of birds that have settled to the north of the Black Sea are uncertain: either they spread there through the Caucasus or came from Ukraine. Recently, single birds were reported from two places in Germany (eBird 2023, GBIF 2023).The current distribution of the SW extends in the west to Austria, in the east to the River Volga in Russia and in the north to the Baltic coast of Poland (Keller et al, 2020; eBird 2023).

In the newly occupied areas, the SW tends to inhabit “human-transformed” open landscapes, such as the parks, gardens, orchards and wooded farmland (Munteanu 1968; Marisova & Butenko 1976; Gorman 2004; Michałczuk & Michałczuk 2016; Figarski & Kajtoch 2018). Although the GW often inhabits more densely wooded areas, including forests proper, this species regularly occurs in “suboptimal habitats” such as semi-open woodlands and “secondary habitats”, and it is here that that two species meet. Interactions between the two species are often agonistic, resulting in territorial disputes, often over potential nesting sites (Munteanu 1968; Michałczuk 2016, Figarski 2017). Nevertheless, there have been, and continue to be, an increasing number of reports of hybrid individuals that have plumage patterns intermediate between the two parental species (Gorman 1999) (Fig. 2). It is known, too, that mixed species pairs (Figarski & Kajtoch 2018) (Table 2) can produce successful broods (Kajtoch & Kusal 2022). Cases of mixed interspecific pairs were often reported in the past (Dudzik & Polakowski 2011); however it is now clear that hybrids of these two species are fertile and, furthermore, there are confirmed cases of successful mixed-hybrid pair broods as well as proof of genome introgression in individuals of a “clear” parental species phenotype (Michałczuk *et al*. 2014; Gurgul *et al*. 2019). Considering the differences in the ecological parameters of these two species, such as preferred nest trees, foraging habitat, wintering ecology and climatic preferences, as well as differences in anatomy (Myczko *et al*. 2020; Pecsics *et al*. 2023), the effect of hybridization on the fitness of resultant birds may be unpredictable. In light of this, the already mentioned competition/aggression and crossbreeding introgression should be considered when studies on SW and GW are conducted in areas where both are present.

**Fig. 2.**
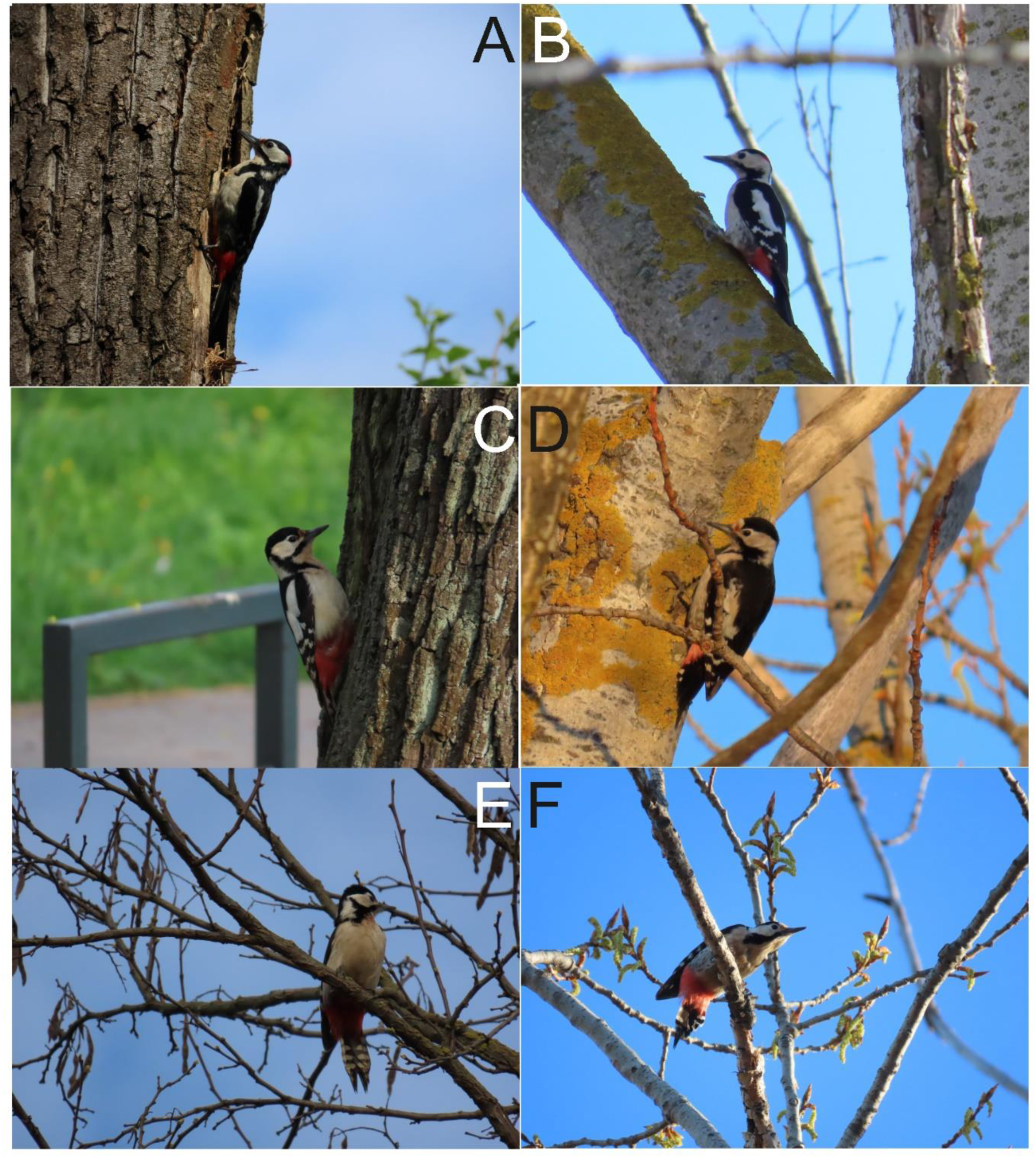
Examples of Great Spotted Woodpecker (A – male, C - female), Syrian Woodpecker (B – male, D - female) and hybrids (E, F).

**Table 2.** Features that aid the identification and distinction of Great Spotted and Syrian Woodpeckers and their hybrids.

| Feature | Great Spotted Woodpecker | Hybrid | Syrian Woodpecker |
| --- | --- | --- | --- |
| Post-auricular stripe | full and wide, extending to the nape | of a various length and width, but never fully connected to the nape | absent or reduced, never extends to the nape |
| Outer tail and undertail feathers | first and second outermost white rectrices forming visible white strip at the tail edges, while undertail overall white with black markings | intermediate coloration between parental species | outermost rectrix is black with white markings, undertail mostly black |
| Undertail coverts coloration | scarlet/deep red | intermediate coloration between parental species | pinkish |
| Red nape patch (in males) | small/narrow | variable | larger (wider) |
| Flanks | clear white | variable | grey stripes, streaked, sometimes pinkish |
| Lores | blackish | variable | whitish |
| White stripes on wings | four-five | variable | three |
| Head shape (jizz) | angular (clear forehead) | variable | pointed (gentle forehead) |
| Bill | stout | variable | narrow, thin |
| Voice | metallic (David Darrell-Lambert, XC458457. Accessible at www.xeno-canto.org/458457) | variable | soft, squeaky (David Darrell-Lambert, XC904299. Accessible at www.xeno-canto.org/904299) |
| Drumming | shorter, faster (David Darrell-Lambert, XC279700. Accessible at www.xeno-canto.org/279700) | - | slower, longer (David Darrell-Lambert, XC904300. Accessible at www.xeno-canto.org/904300) |
| Feeding sites | most tree species, deciduous at breeding season, conifers at winter | same as parental species | softwood trees and fruit trees in lightly wooded country |

The aims of this paper are to:

1. To summarize the current state of knowledge on research dealing with the distribution, demography, biology and ecology of SW and GW.
2. To assess how many studies on GW and SW were undertaken in the sympatric range of both species but which did not consider both species.
3. To summarize how many studies have dealt with the co-occurrence, competition or hybridization of these two species.
4. To find how many studies considered both species (regardless of the goal of study) and what patterns were observed.
5. To estimate which biases could result in studies on these species when only one was studied despite both being present in the study area.
6. To recommend goals for future research on these two species.
7. To highlight the need to consider both species in monitoring and/or mapping.

## 2. Methods

### 2.1. Literature search

To review the current state of knowledge and problematic issues described above, a search of the scientific literature, using the Web of Science (WoS) and Scopus electronic databases, was conducted in November 2023, using the following key words: “*Dendrocopos major*” and “*Dendrocopos syriacus*”. The downloaded metadata was organised into a single sheet having information on the title, authors, year of publication, keywords and journal. Each text was briefly examined based on its title and abstract, to provide a short classification in the form of keywords or to eliminate the publications that are not connected to the research goals (e.g. there were cases when studies found dealt with other species, such as beetles, and only briefly mention GW and/or SW). Topics which were not directly connected to our study were excluded from the analysis and the reason for the rejection was noted. First, articles obtained from WoS by inserting the key words “*Dendrocopos major*” were examined, then articles obtained from the same database using the key words “*Dendrocopos syriacus*.” Subsequently, the same method was used to retrieve papers from Scopus. Finally, duplicates from both databases were deleted (Fig. 3).

**Fig. 3.**
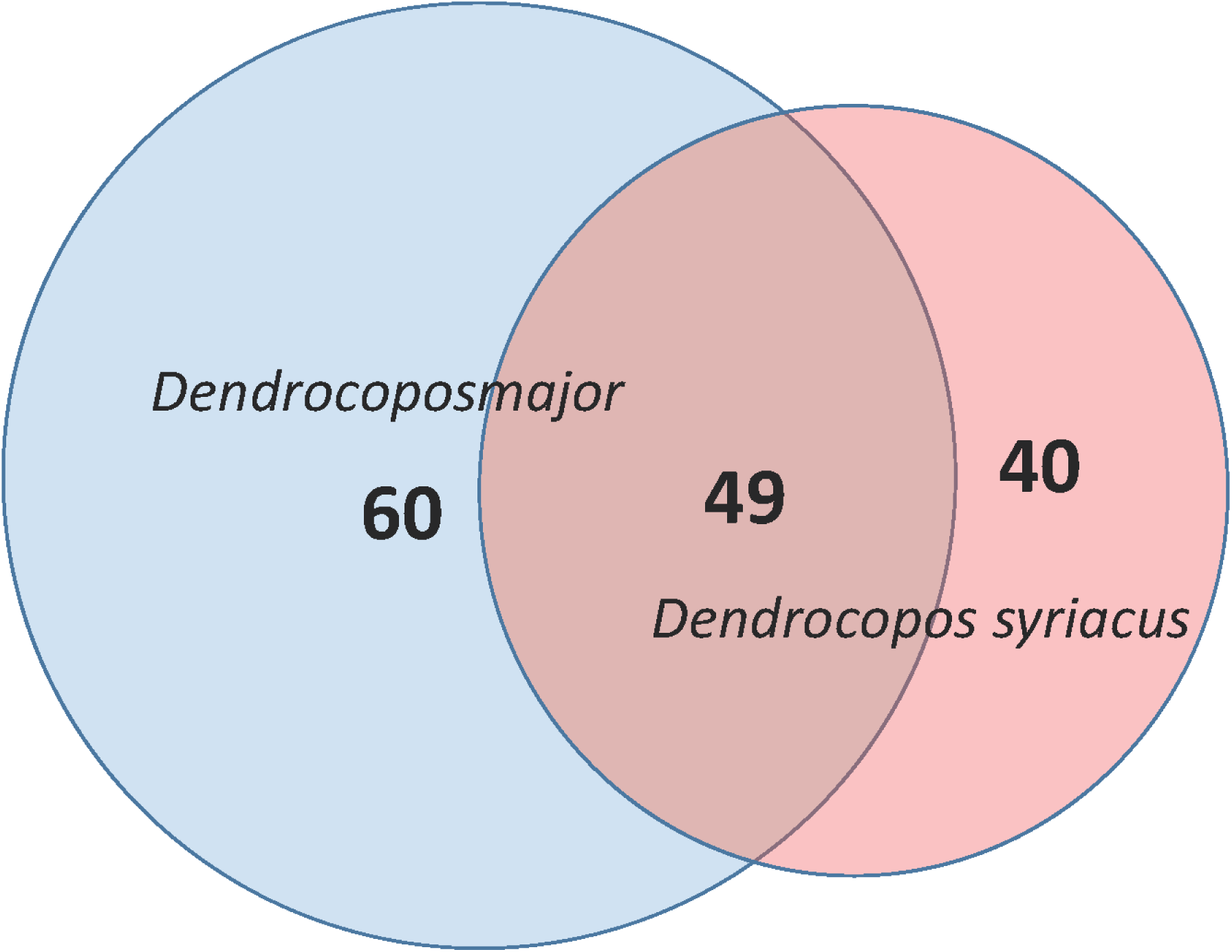
Venn diagram showing number of studies conducted on great spotted and Syrian woodpeckers (including studies on both of them).

### 2.2. *Data* analyses

After filtering the obtained data, we selected only papers originating from the sympatric range of both species (Iran, Caucasus, Turkey, Balkans, Carpathian Basin, Poland, Belarus, Ukraine, Russia). In this pool, all of the scientific works found were tagged according to the content (Table S1), the country where the research was conducted, or from where the woodpeckers originated, and the species with which the article dealt with (Fig. 3).

Due to the nature of much of the available data being descriptive, the possibility of further analyses was extremely limited. Therefore, most of the questions of this study were answered by referring to particular studies and comparing their data and results.

## 3. Research description and summary

### 3.1. Basic summary of the search results

From the WoS database 1017 texts were obtained by searching with “*Dendrocopos major*” keywords, and 174 texts by “*Dendrocopos syriacus*”. From the Scopus database 706 texts were obtained using “*Dendrocopos major*” and 106 using “*Dendrocopos syriacus*”. To identify duplicates in WoS “*Dendrocopos major*” search results were used as a standpoint. Overall, this resulted in 10 articles from GW WoS, 36 from DS WoS, 152 from GW Scopus, and 49 from DS Scopus. However, the last search produced 7 duplicates in Scopus (Fig 4). 457 articles from WoS (6 data study, 4 phylogeography, 5 urban damage, 5 anomaly, 2 biomechanics, 12 methodology, 178 not about species, 4 palaeontology, 17 pest control, 63 faunistic, 35 general ecology, 20 photographic records, 16 parasitology, 9 physiology , 7 ethology, 5 ringing, 2 cytology, 4 anatomy, 2 figure, 4 morphology, 6 epidemiology, 4 nature conservation, 3 microbiology, 2 histology, 2 sociology, 1 general book, 1 bioinformatics, 1 urban management, 3 ecotoxicology, 33 other) and 583 articles from Scopus (335 not connected to species, 6 biotechnology, 32 nature conservation, 42 epidemiology, 74 general ecology, 5 pest control, 11 methodology, 11 microbiology, 13 other, 8 natural history, 1 phylogeography, 1 agriculture, 3 physiology, 1 general evolution, 1 forestry, 2 taxonomy, 5 ringing, 13 faunistic, 2 informatics, 1 morphology, 4 ethology, 2 subspecies, 6 urban damage, 1 anatomy, 1 molecular biology) were excluded as they that did not specifically relate to our study. For 294 articles we were not able to confirm a source, thus these were also excluded from our analysis. After filtering out all the duplicates (247) - topics not relevant to our research (1040) and articles that were not available (294) - we obtained 424 articles of which 149 were from the sympatric range of the two species (Table S1, Fig. 4). There is an substantial increase in number of studies on these two woodpeckers with particular accelerating number of articles after 2015 (Fig. 5).

**Fig. 4.**
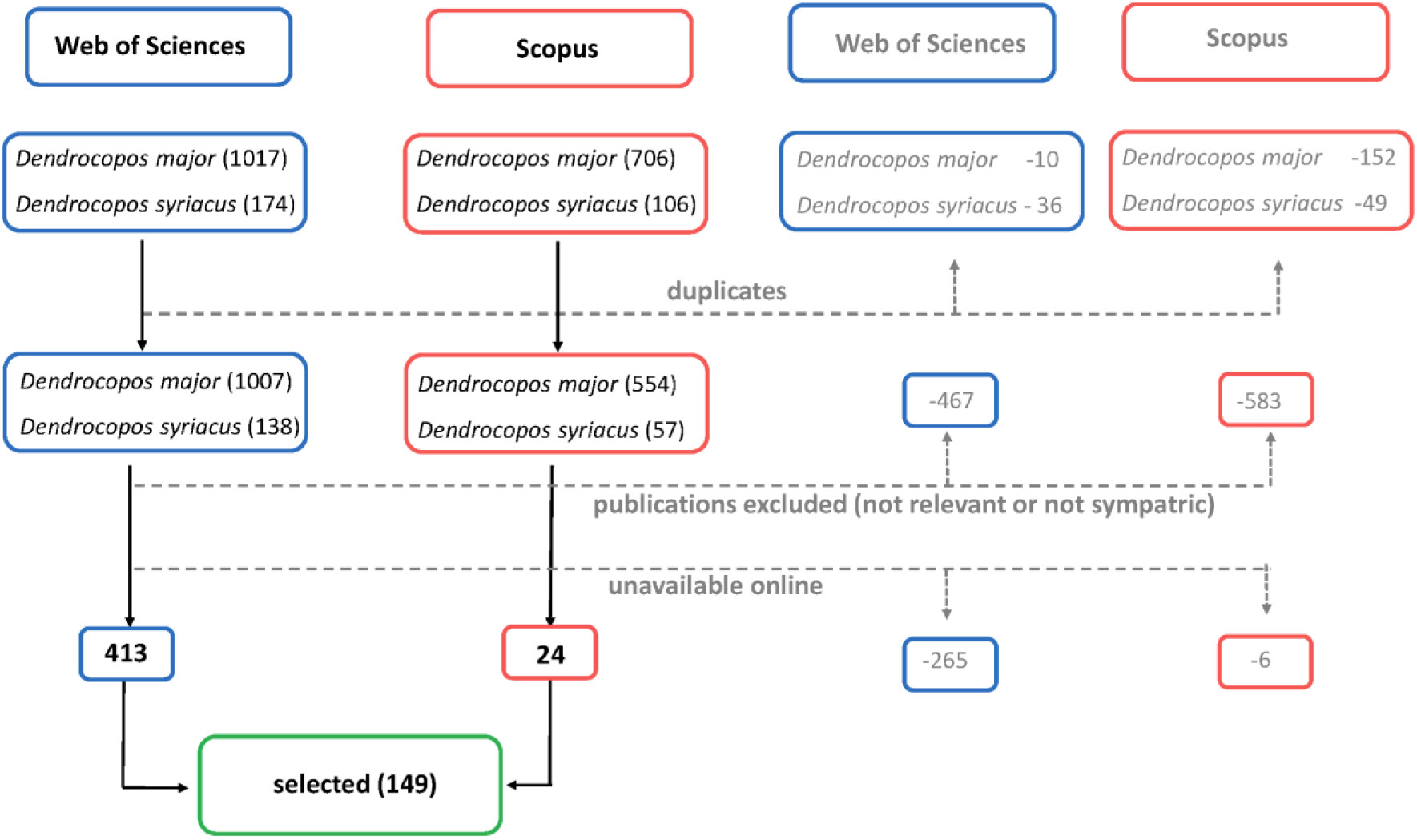
PRISMA (Preferred reporting items for systematic reviews and meta-analyses) workflow presenting search and selection of relevant articles in Web of Sciences and Scopus databases for purposes of the review summary.

**Fig. 5.**
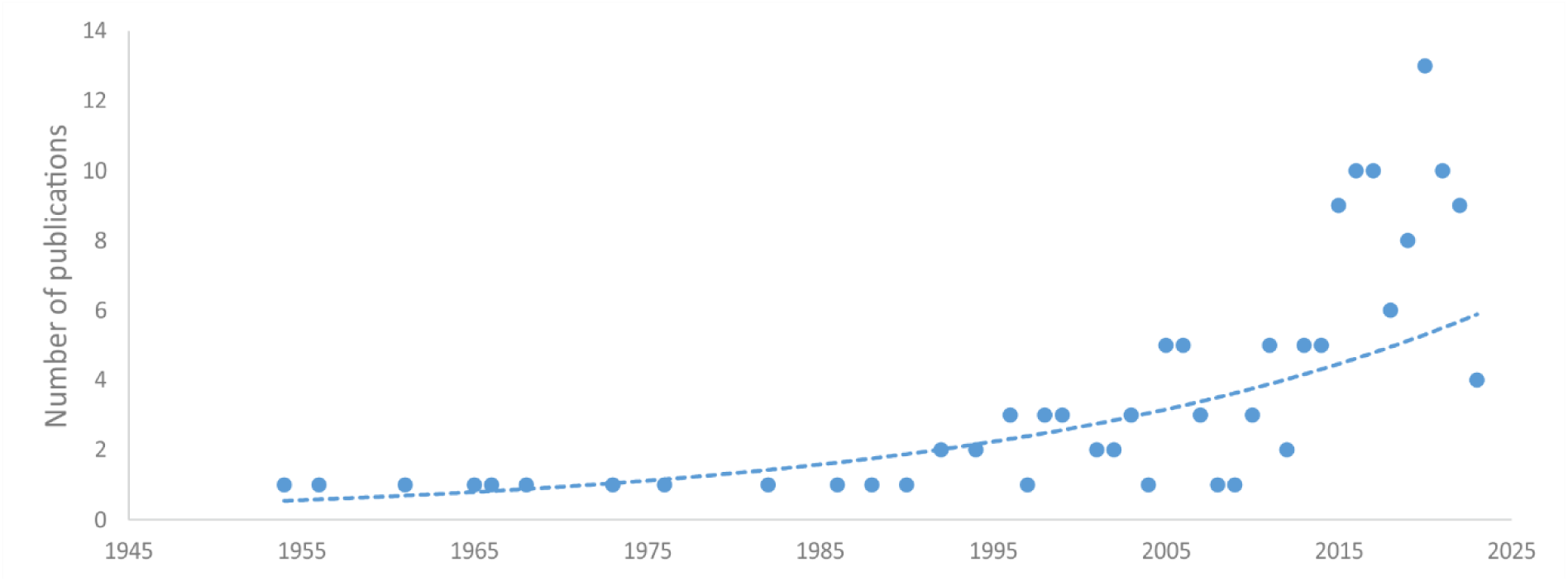
Progress in studies on Great Spotted and Syrian Woodpeckers in areas where they are sympatric.

### 3.2. Geographic span of research

The majority of articles that deal with these two species together are from Poland (Fig 6), mostly from the south-east and east, where studies on SW and their hybrids are actively conducted. A substantial number of texts also come from the Pannonian region: most concerning Hungarian birds though three publications were found Romania. Numerous texts related to birds in Russia, but all were either short notes on observations of “unusual” behaviour or records of SW from local areas. There were, however, two summaries of the range expansion through the Russian by SW and three articles focusing on GW ecology. In Slovakia, only two publications mentioned SW breeding and one report on an unidentified “albino”, whereas in Czechia there was one study on population dynamics in a local bird community and one report of a hybrid. The other ten papers from these regions concerned GW ecology and demography. Literature from the Pontic region (eastern and southern Romania, Moldova, Ukraine not including the Carpathians) contained six old (1950-1970) papers describing SW ecology, range expansion and distribution, two censuses, one recent description of the settling of SW in the Ukrainian steppe zone and one paper about GW breeding ecology from Romania. Two publications from Belarus were found, one concerning GW trophic ecology and a note on SW. The search result from the Western Balkans (Slovenia, Croatia, Bosnia and Herzegovina, Serbia) showed three records of SW breeding and one note on GW. Publications from the area of original distribution of SW in the Middle East are lacking. We found just two papers dealing with GW ecology and four publications on SW ecology and breeding biology from Iran and one paper that mentioned both species in oak forest-pasture in Turkey. The entire range of the SW distribution was mentioned in Michalczuk (2014) and the expansion and European distribution of this species discussed in Gorman (1996).

**Fig. 6.**
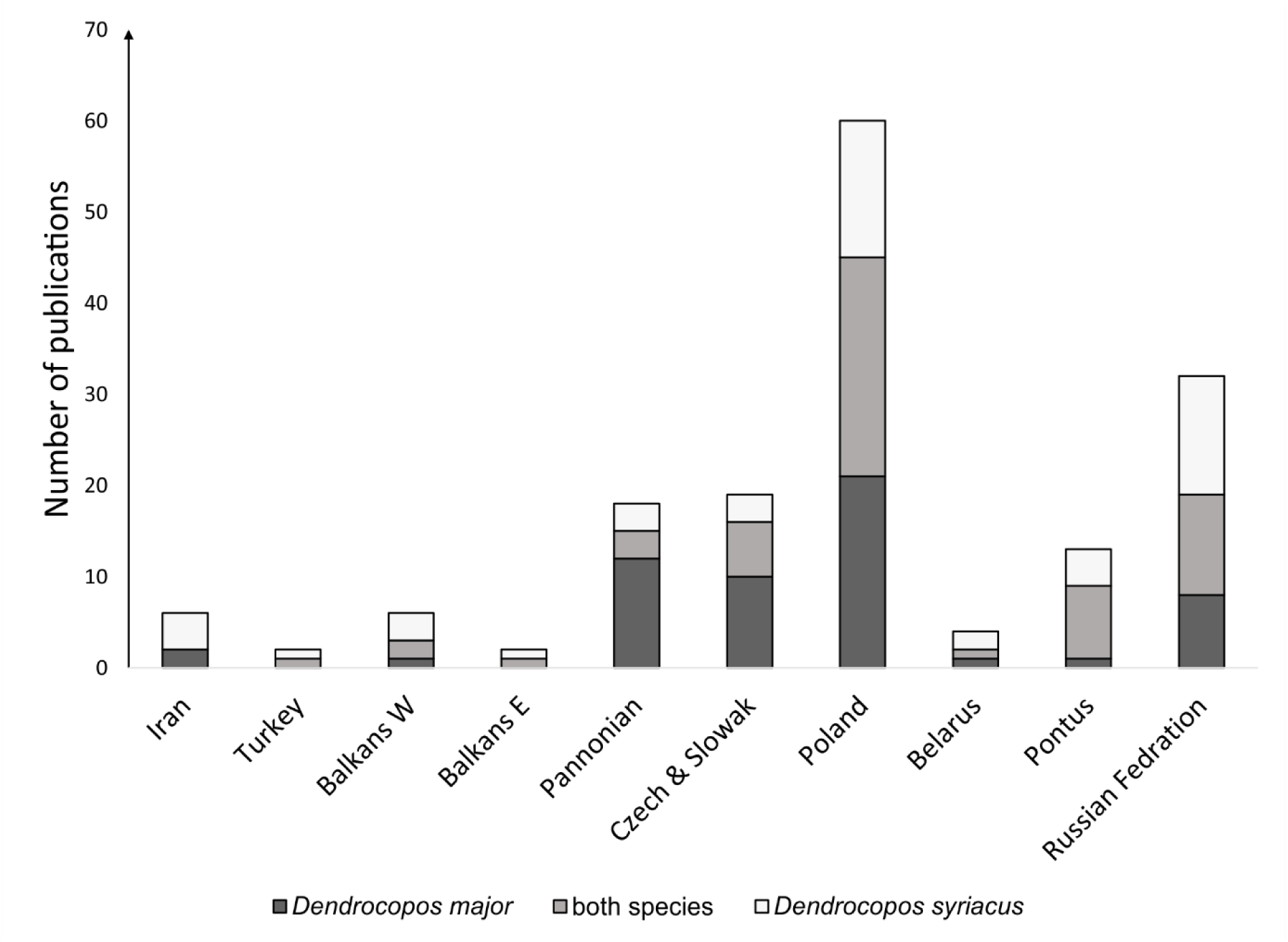
Distribution of studies on Syrian and Great Spotted Woodpeckers over countries and regions in Europe and the Middle East.

### 3.3. Birds’ morphology

#### 3.3.1. GW studies

Among the literature examined, two studies concerned atypical coloration: one about unusual GW individuals (Olszewski 2007), and another on an unidentified “albino” (Karlik & Vel’ky 2007). Thirteen studies compared the plumages of both species: the main topic in three of these was plumage differences and identification (Horvath 1961; Gorman 1996, 1999), while the other ten only mentioned differences and hybrid plumage (Matousek 1954; Marisova 1965; Dudzik & Polakowski 2011). Two studies dealt with the bill and cranial morphometry of these two species together with those of other European woodpeckers (Myczko et al. 2020; Pecsics et al. 2023). Vocalisation differences between GW and SW were mentioned in seven publications (Munteanu 1968; Krotoski, Karnas & Przyszlak 1986; Dudzik & Polakowski 2011).

#### 3.3.2. SW studies

In addition to the publications mentioned above, there was an additional article in which SW identification was mentioned (Matousek 1954). In an interesting case from Russia, local researchers decided to re-examine their photos of GW from previous years and discovered that some of the birds were actually SW, although they did not find any hybrids amongst them (Valova & Fionina 2018).

### 3.4. Methods used for woodpecker studies

The majority of publications analysed were field surveys (121) (Table 3). Playback stimulation was only used in twenty-six of these (3 from Romania (Dorrestejin et al. 2013; Damoc et al. 2014; Domokos & Cristea 2014), one from Iran (Mohamadian, Shafaeipour & Fathinia 2019) and the rest from Poland). Publications in which field research was not conducted included books and articles (Gorman 1996, 1999, 2015), studies on museum specimens (Myczko et al. 2020; Pecsics et al. 2023), dead specimens (Hanak, Rumler & Vermouzek 2003), a mathematical model (Pavlik 1994), summaries based on other publications (Marisova 1965; Pasinelli 2006), nest cards (Hebda & Szewczyk 2005; Hebda 2009) and ringing data (Ciach & Fröhlich 2013). In addition, there was one molecular study on phylogeographic patterns of GW (Zink, Drovetski & Rohwer 2003), and two molecular studies on both species and their hybrids (Michalczuk et al. 2014; Gurgul et al. 2019). One study dealt with Hungarian SW and GW phylogenetics based on plumage features (Horvath 1961). Another study of GW population dynamics was based on ringing data and satellite photos of landscapes (Onodi & Csorgo 2014). For SW there were also five summaries on range expansion (Keve 1956, Sokolov 2020, Strautman 2023).

**Table 3.** Summary of goals in studies on Great Spotted and Syrian Woodpeckers.

| Category | great-spotted<br>woodpecker | both<br>woodpeckers | Syrian<br>woodpecker |
| --- | --- | --- | --- |
| field observation | 54 | 35 | 32 |
| playback used | 6 | 13 | 7 |
| molecular | 1 | 3 | 0 |
| conservation | 13 | 7 | 3 |
| ecology | 53 | 31 | 27 |
| habitat requirements | 44 | 33 | 27 |
| foraging/diet | 28 | 13 | 10 |
| nesting | 15 | 14 | 17 |
| reproductive biology | 9 | 13 | 13 |
| density/distribution | 28 | 22 | 16 |
| demography | 10 | 15 | 17 |
| plumage | 1 | 14 | 2 |
| dimorphism | 2 | 5 | 0 |
| phylogenetics | 1 | 2 | 0 |
| population genetics/phylogeography | 1 | 1 | 0 |
| hybridization | 0 | 19 | 3 |
| competition/territorialism intraspecific | 5 | 5 | 1 |
| competition/territorialism interspecific | 7 | 12 | 4 |
| breeding | 44 | 34 | 28 |
| non-breeding | 18 | 17 | 10 |
| urban | 6 | 25 | 18 |
| rural | 12 | 20 | 23 |
| forest natural | 39 | 6 | 3 |
| forest plantation | 8 | 0 | 0 |
| riparian | 10 | 6 | 5 |
| summary | 415 | 365 | 266 |
|  | 780 |  | 631 |

### 3.5. Habitats

#### 3.5.1. GW studies

The majority of studies on GW were conducted in forests (39), mostly “semi-natural” but two were from plantations (Walankiewicz et al. 2011; Czeszczewik et al. 2013) and one used a traditional rural landscape for comparison (Dorresteijn et al. 2013). One compared populations in forest, swamp and rural areas (Ivanchev 2017), one studied the differences between natural, riparian and plantations (Domokos & Cristea 2014), and two publications covered all of these habitat categories (Hebda 2009). Eight studies on GW were conducted solely in rural landscapes (Merzlikin & Sheverdyukowa 2005; Mazgajski & Rejt 2006; Krnjeta 2019), two in plantations (Kebrle 2021; Velova, Vele & Horak 2021), four in urban areas (Sychra 2003; Kocian et al. 2003; Melnikov, Belachenko & Belachenko 2017) and three in riparian forests (Onodi & Csoergo 2014; Onodi & Winkler 2016; Onodi et al. 2021). Others compared riparian forest to plantation (Kosinski, Ksit & Winiecki 2006).

#### 3.5.2. SW studies

Almost all studies on SW were conducted in urban (18) or rural (23) areas, except for three studies from Iran, where birds are present in “natural” steppe-forests (Aghanajafizadeh et al. 2011; Mohamadian, Shafaeipour & Fathinia 2019; Shafaeipour, Fathinia & Pasinelli 2020). Five studies also included riparian forests (Havlin & Havlinova 1966; Gasik 2010; Belyachenko & Melnikov 2018).

#### 3.5.3. GW & SW studies

Studies which included both species were mainly conducted in urban (25) or rural areas (20). Some studies included “natural” woodlands (6) (Hebda & Szewczyk 2005; Michałczuk & Michałczuk 2016; Askeyev et al. 2022) and riparian forests (6) (Ciosek & Tomiałojc 1972; Hubalek 1999; Kajtoch & Figarski 2017) along with rural ones. No papers were found that dealt with both species in forest plantations.

### 3.6. Research goals / topics

More than half of the publications examined were connected to the ecology of the studied species (108). Many included such information as diet or feeding behaviour, habitat requirements and breeding biology (Table 3).

The majority of studies were conducted during the spring breeding season (106) with fewer outside that period (45). Most of the autumn-winter studies dealt with GW, and mainly concerned winter diet (10) or year-round diet (7), and almost all of these were ecology focused. In just nine out of all of the winter papers, the subject of study was SW biology, and fifteen texts were related solely to winter records.

#### 3.6.1. GW studies

Publications about GW mainly dealt with its ecology (53), habitat requirements (44) or diet and/or feeding behaviour (28). Fifteen studies described nests and nine had data on breeding biology. Twenty-eight studies focused on population density and distribution, and nine on demography. One study focused on phylogenetic patterns of GW across Eurasia (Zink, Drovetski & Rohwer 2002). Seven studies mentioned interspecific competition for territory (Wesolowski 1990; Pavlik 1994), food resources in winter and competition between males and females (Ivanchev 2017; Melnikov, Belyachenko & Belyachenko 2017). Five studies included information on GW competition with Lesser Spotted Woodpeckers (*Dryobates minor*) and Middle Spotted Woodpeckers (*Dendrocoptes medius*) (Pavlik 1992; Stanski et al. 2020; Onodi et al. 2021).

#### 3.6.2. SW studies

The majority of papers on SW focused on the ecology of the species (27), with most including data on habitat requirements (24) and some on diet and foraging behaviour (9).

Information on SW habitat requirements was also included in two faunistic records, and one record included notes on food items. Seventeen publications had information on nest characteristics and thirteen about breeding biology. Sixteen texts mentioned population density and distribution and 17 demography. Two faunistic reports mentioned the possibility of interbreeding with GW (Goroshko & Kusenkov 2004; Gasik 2010). Two field observations documented territorial/agonistic behaviour towards GW and vice versa (Goroshko & Kusenkov 2004; Gizatulin & Khokhlov 2017). One brief summary contained statements on agonistic behaviour towards GW and competition for territories (Mityai 2022).

#### 3.6.3. GW & SW studies

Works where both species were treated together included thirty-one ecological studies. Such research mainly describes habitat requirements (Michałczuk & Michałczuk 2016; Figarski & Kajtoch 2018; Michałczuk et al. 2018), with some on population dynamics (Listopadski 2016; Askeyev et al. 2020; Kajtoch & Kusal 2023) and diet (Marisova 1965; Winkler 1973; Czyż & Celiński 2012). Some studies detailed mixed pairs (Bacani 1998; Dudzik & Polakowski 2011; Figarski & Kajtoch 2018) or hybrids (Krotoski, Lontkowski & Przyszlak 1986; Harrop 2005, Kajtoch & Kusal 2022). Some reports (Lorek & Durczynska 1992; Ganitsky 2018; Moskvichev 2020) and summaries (Keve 1956; Zavialov, Tabachishin & Mosolova 2008; Sokolov 2020) focused on SW range expansion but also mentioned GW. In most studies population densities of these woodpeckers were presented (Hubálek 1999; Michałczuk & Michałczuk 2016; Kajtoch & Figarski 2017). Three molecular studies cited both species and their hybrid genomes (Michałczuk et al. 2014; Gurgul et al. 2019; Kajtoch & Kusal 2022).

### 3.7. Research dealing with both species

#### 3.7.1. GW and SW woodpeckers’ ecology in rural landscapes

GW has seldom been studied in rural landscapes but there are studies from habitat mosaics in Hungary (Onodi & Csorgo 2012) and an agricultural landscape in western Poland, in an area where SW is scarce (Jermaczek 2022).

Both species were assessed in Czechia as a part of study on the seasonal changes in bird communities in a managed lowland riverine ecosystem (Hubálek 1999). A study on woodpeckers in riparian forests in Romania (Domokos & Cristea 2014), included GW but not SW.

The majority of studies on rural populations of SW have been conducted on breeding pairs in SE Poland (Michałczuk & Michałczuk 2016; Kajtoch & Figarski 2017; Michałczuk & Michałczuk 2020…). Michałczuk & Michałczuk (2016, 2020), which are the only detailed source of data for SW breeding in orchards. These studies, undertaken within the core Polish population, show that this species seems to favour old orchards for breeding. In Michalczuk & Michałczuk (2020) habitat requirements were described and in Michalczuk (2016) details about the reproductive biology of SW were given. Most of the conclusions in these works were later confirmed by Kajtoch & Figarski (2017) and Kajtoch & Figarski (2018) in another area in southern Poland. Orchards can be categorized into three types: old and traditionally cultivated, intensively farmed with small trees, and overgrown abandoned orchards. SW prefers the first type. Further studies proved that the decline of these populations was due to the loss of traditional horticultural practises (Michalczuk 2015; Kajtoch 2023).

Some of the above mentioned papers considered the question of the coexistence of SW and GW. At the beginning of the 21^st^ century in SE Poland, Michalczuk (2016) showed the spatial segregation of these species. GW tended to occupy remnants of forests found in agricultural mosaics and therefore the two species rarely met. This situation seemed to change after 2015, when GW increased in number and also started to breed in typical rural woodlands. The increase in GWs in rural landscape likely accelerated the decline of SWs (territory replacement) but simultaneously, it enabled an increase in interspecific mating (Figarski & Kajtoch 2018).

#### 3.7.2. Determinants of GW and SW occurrence in urban woodlands

In urban areas GW was mostly studied in wooded areas situated within cities, therefore these populations should not really be treated as truly “urban.” For example, Mazgajski (1997, 1998, 2000, 2006) assessed populations of GW in urban woodlands in Warsaw (sometimes described as “parks”), but no information on any co-occurring SW were included. Kopij & Holga (2008) detailed the occurrence of woodpeckers in woodslands in the city of Wroclaw, but again no data on SW were reported (although this species breeds there).

A well-studied urban population of various woodpecker species is found in Krakow in southern Poland. First, Ciach & Fröhlich (2013a, b) described the occurrence there of SW and suggested that this species benefits from air pollution, as it excavates nest holes in trees that have been weakened by pollution in city centre. However, this study was based on occasional observations collected over an extended period for bird atlas purposes and neglected sites in the suburbs. Later, the same authors (Fröhlich & Ciach 2020) described the relationship of woodpeckers (including GW and SW) with the presence of deadwood in cities. Figarski & Kajtoch (2018) investigated urban populations in central Poland and revealed niche shifts between SW and GW in urban areas, with only partial overlapping of use of habitats.

According to the evaluation of the available literature, the main determinant of SW occurrence in urban areas is the presence of large, old, weak and deceased softwood trees such as poplars *Populus* spp., willows *Salix* spp., black locust *Robinia pseudoacacia* (Ciach & Fröhlich 2013a; Gorman 2020), which are used as both foraging and nesting sites. The scattered, open, distribution of trees in breeding areas is also considered important (Ciach & Fröhlich 2013b). Kajtoch & Figarski (2017) also emphasised the importance of old gardens in urban landscapes. In cities, GW was found to benefit from a higher volume of wood stands and a higher number of mature trees (Fröhlich & Ciach 2013). Both species benefit when old trees and deadwood are present in their habitat (Figarski & Kajtoch 2018; Fröhlich & Ciach 2020; Froehlich, Hawryło & Ciach 2022).

#### 3.7.3. The ethology of sympatric SWs and GWs

Only one study was found that focused on the behaviour of sympatric SW and GW in urban environments. Figarski (2018) monitored the behaviour of both species, including observing the sexes separately, throughout the year. His research showed differences in vocal activity with signs of a higher aggressiveness from SW females when they met GW females. Moreover, this study showed that territories are not constant over a year, although adult birds usually do not move far from their breeding sites. In addition, some information on these species’ interactions was also gathered from several brief ornithological notes where authors shared their field observations (Zavyalow 1996; Melnikov 2015; Listopadski 2016) and from articles where the main subject was not ethology (Winkler 1973). Overall, authors generally reported a strong territorial aggression that is shared between GW and SW and manifests itself in interspecific interactions.

#### 3.7.4. Mixed pairs and hybridization

The first records of hybrids between SW and GW were from 1950s and 1960s from the Carpathian Basin (Austria and Hungary) (Keve 1956; Winkler 1971; Gorman 1997; Michałczuk 2014). For a long time, however, there were only occasional reports of mixed pairs or suspected hybrids in Central Europe, which suggested that this phenomenon was rare and hence did not have an impact on the reproduction of the two species. Whether hybrids were fertile was unknown and it was surmised that if indeed they were, they would be “less fertile” than “pure” birds. In 2011 Dudzik and Polakowski published a local paper on hybridization between SW and GW in Poland where they reported several cases of mixed pairs and hybrids along with a summary of their plumages and occurrence. That work changed the approach to the question of hybridization between these two woodpeckers.

Further studies were undertaken in Poland, particularly in Krakow, where an increasing number of mixed pairs and hybrids have been detected since 2014 (Figarski & Kajtoch 2018, Gurgul et al. 2019). In this Polish population, not only mixed pairs composed of “pure” individuals of both species, but also pairs of hybrids (backcrosses) and even a cases of “double hybrid” pairs with successful broods, have all been confirmed (Kajtoch & Kusal 2022).

## 4. Discussion

The expansion of SW into Europe is considered to have been a natural process as the species spread from Anatolia into the Balkans without any direct human help (Glutz von Blotzheim & Bauer 1980; Michałczuk 2014; Strautman 2023). Nevertheless, its expansion was most likely enabled by indirect anthropogenic change - the development of rural woodlands, mostly orchards, which, however, had a much older history preceding expansion of SW (Listopadsky 2016). It is likely that some other factors also forced or enabled the species to spread e.g. an increase in abundance in its native range, or a decrease in required habitats for nesting or foraging. Climate change was probably not responsible for the initial expansion, as it began before the acceleration of global warming began (Bradley 2000).

Nonetheless, in recent decades, the demography and distribution of SW might have been affected by climatic factors as its range expansion to the north-west, where the climate is probably too humid in winter, ceased. Rather, the movement of SW shifted to the north-east where the continental climate is ostensibly more favourable for this “forest-steppe” bird (Askeyev 2022). Nevertheless, the inclusion of an additional species into the European woodpecker assemblage most likely affected the distribution, ecology and ethology of the already existing members, particularly the most closely related species, GW.

Indeed, the subsequent sympatry of SW and GW has driven various reactions in both species. Unfortunately, very few studies investigate and describe these interactions.

### 4.1. Data availability

It seems that SW, in particular the issue of its hybridization with GW, is most actively studied int SE Poland. Most publications on this species come from that area owing to the intensive research of groups such as those lead by Dr J. Michalczuk in the Roztochchia upland, Dr. M. Ciach in Kraków city and Dr T. Figarski in several cities in Poland (Ciach & Frohlich 2013; Michałczuk & Michałczuk 2016; Kajtoch & Figarski 2018). Little information on SW exists from the area of its original distribution in the Middle East, Iran and Turkey) and from the region where its range expansion was first noted (the Balkans), while hybridization is not even mentioned. One article from the Taurus Mountains in Turkey reports that both species were present in wooded pasture, but no information on their interactions were presented (Bergner et al. 2016). The few publications that exist from Ukraine indicate a lack of inclusion in open electronic data-bases rather than an actual absence of observations and research on SW. Still, this statement could also apply to some other countries.

### 4.2. Co-occurrence

Most of the available published resources on SW and GW simply describe their respective occurrences, usually as part of wider studies on the whole woodpecker or total avifauna assemblages. Therefore, data from these papers are generally of limited use and seldom lead to important findings that extend our knowledge of eco-ethological interactions between the two species. Some are simply ornithological notes of unusual behaviour (Melnikov 2015) or the confirmation of SW as a new breeding species for a particular region (Lorek & Durczynska 1992). Often, studies and notes describe the distribution and abundance of SW and GW in rural or urban landscapes where they occur together (Ciosek & Tomialojc 1982; Micałczuk & Michałczuk 2022). This review intentionally omits studies on forest populations of GW, as SW is known to avoid such closed habitats. In short, SW tends to dominate in number in the open rural and urban wooded landscapes of Central, East and SE Europe, while GW prefers to breed in forests proper, including forests on the edges of cities (Michałczuk & Michałczuk 2016; Figarski & Kajtoch 2018). Reports from the time of the SW expansion, often emphasised that this species replaced GW in anthropogenic habitats (Munteanu 1968; Listopadski 2016). However, more recently, some papers have reported a reverse phenomenon - a decrease of SW populations in rural and urban landscapes with the simultaneous expansion of GW (Kajtoch & Kusal 2023). This concerns mostly populations in southern Poland, although some reports suggest a similar trend all over Central Europe.

Unfortunately, reliable trends of population dynamics are known only for GW, which is mainly due to this species being included in the monitoring of common breeding birds in EU countries (https://pecbms.info/). Such monitoring suggests a substantial increase in GW numbers, which can explain its colonisation (or recolonisation) into urban and rural woodlands, as a result of over density in its core forest populations. The recent decline of SW in some countries (notably Poland, but also Romania, and there are reports of a decline in numbers also in the Levant), is a trend that may be important for the conservation of this species which is overall rather rare within the European Union (e.g. annexed in the Bird Directive) (https://datazone.birdlife.org/species/factsheet/syrian-woodpecker-dendrocopos-syriacus/details).

### 4.3. Interactions

#### 4.3.1. Aggressiveness

Some studies have described direct interactions between SW and GW (Winkler 1973; Krotoski et al. 1986). The majority of these interactions are competitive or aggressive in nature. However, there is no unambiguous evidence to prove whether one species is dominant over the other (Gorman 2004). Individuals of both species have been observed to be the more aggressive in disputes with their congener (Lorek & Durczynska 1992; Figarski 2017). GW is known to be aggressive to other woodpeckers that share its habitats, particularly to Middle Spotted, White-backed and Three-toed Woodpeckers *Picoides tridactylus* (Friedman 1993; Nowak 2003). Its behaviour towards other woodpecker species is usually limited to chasing them away. However, territorial aggression between GW and SW is a complex phenomenon as both taxa show interspecific aggressiveness, using characteristic vocalizations and displays. It seems that SW and GW partition a common environment as their habitat niches only partially overlap and there thus remains some space for co-existence (Michałczuk & Michałczuk; Figarski & Kajtoch 2018). Likely niche displacement depends on the densities of both species and in sites where one is dominant, the other species can be forced to occupy less favourable sites. Indeed, this is a common phenomenon among close relatives in birds (DeBach 1996).

#### 4.3.2. Cooperation

Surprisingly, it seems that there is only one report that details a positive interaction between GW and SW, namely cooperative breeding. Melnikov (2015) reported that a lone GW female “helped” to raise the nestlings of a SW pair in Saratov oblast (Russia). It is likely that this behaviour was caused by the loss of the brood (and/or mate) of the GW female, and instinct drove this female to provide food to the SW brood.

### 4.4. Mating and hybridization

#### 4.4.1. Polyandry

GW and SW were traditionally considered to be monogamous species, although Kotaka (1998) proved that GW could be polyandrous. There are no published reports on this subject that involve SW, although personal observations of the authors suggest that these two species could mate with members of the other species in a way of extra-pair copulations. In 2022 a hybrid female (paired with a hybrid male) in Krakow was seen to copulate with a GW male from an adjacent territory (that was paired with GW female). In 2024 copulation between a SW male and a GW female, from pairs that bred adjacently, was observed in Proszowice, southern Poland.

#### 4.4.2. Mixed pairs

Recent studies have proved that mixed pairs regularly occur in sympatric populations in SE Poland (Figarski & Kajtoch 2018). Initially, the share of such pairs was estimated to be around 5%, but that share seems to have increased in recent years (between 2014-2015 and 2022-2023) as more mixed pairs (including pairs of hybrids) have been seen in Kraków (currently up to 20%, unpublished). Observations suggest that most mixed pairs are composed of a SW female and a GW male, or a hybrid female and a GW male (Dudzik & Polakowski 2011; Figarski & Kajtoch 2018). This intimates a degree of assortative mating, although the reasons for this remain uncertain. They may be biological/ethological or demographic, related to the densities of potential mates in a given population.

#### 4.4.3. Hybridization

SW and GW hybrids have been reported for decades, but only sporadically (Hanak, Rumler & Vermouzek 2003; Harrop 2005). First, such birds were mainly found in Hungary, but later mostly in Poland. Indeed, in Poland, field and genetic studies undertaken in 2012- 2024 proved the occurrence of hybrids on a much larger scale than was suspected.

Approximately 4% of birds observed in some sympatric populations (particularly urban) were identified as hybrids and this number increased to 7% for birds verified in the hand (e.g. dead or captured individuals (Figarski & Kajtoch 2018). Implementation of genetic markers (mitochondrial DNA, introns, microsatellites and single nucleotide polymorphisms) revealed that 20% of birds had some introgression genes from both species (Michałczuk et al. 2014).

Finally, it was proved that two hybrids (so-called “backcrosses”) are also able to mate and successfully produce offspring (Kajtoch & Kusal 2022). All of the above strongly suggests that the hybridization of SW and GW is not an uncommon phenomenon, particularly at the edges of the SW range. As already mentioned, examination of photographs from the Russian literature revealed that some birds previously classified as pure SW where actually hybrids (Melnikov 2015; Moskvichev 2020). The first publication about SW in Russia where the authors accounted for the possibility of hybridization, and hence asked experts to evaluate the “purity” of the individuals in photographs, was written by Valova & Fionina (2018). The same authors reported that during censuses between 2010-2015 they did not consider the presence of SW at all, and simply recorded them as GW. Only later, when revisiting old materials, did they discover that some individuals were in fact SW. Reports on eBird have also mentioned cases of possible hybrids in Belarus, Croatia and even Turkey. Thus, hybrid SW x GW could even be present with the established, core range of SW.

### 4.5. Consequences of SW and GW interactions for scientific research

Both competition/aggressiveness and mating/hybridization in SW and GW have probably had an impact on the biology, ecology and ethology of these species. First of all, the co-occurrence of these two species presumably results in some displacement as their ecological niches overlap. It seems that the expansion of one species into a range of another (SW into GW in the past, and currently GW expansion into urban/rural woodlands occupied by SW) leads to territory shifts or even displacement (Listopadski 2016; Michałczuk & Michałczuk 2016). There is no definitive proof that either SW or GW is dominant with respect to individuals, as both actively defend their territories, and a likely shift or replacement is either random (depending on the level of aggressiveness of individual birds) or depend on local densities (dominant birds occupy most of the territories). There are some suggestions that SW females could be more aggressive than GW females (Figarski 2017), and this could explain cases of polyandry and interspecific mating. At the moment, little is known about differences in the reproduction between allo- and sympatric populations of GW and SW. That is, if sympatric one or both of these species have a lower breeding success due to competition for resources (space, food). It is likely that there is no defining “rule” and differences cannot be observed on the level of individual pairs but on the level of whole populations (the overall decrease in the number of one species). The differences in morphological adaptations to excavation and food acquisition between the two species (Myczko et al. 2020; Pecsics et al. 2023) make assessing hybrid fitness challenging, as almost no data connected to this subject is available. Unfortunately, most of the studies undertaken on woodpeckers in rural or urban landscapes focus on either GW or SW, neglecting the role of the sibling taxa (Fröhlich & Ciach 2013; Ónodi & Winkler 2016). Some research considers both species as a part of topics executed on the whole woodpecker assemblage but ignore the presence of mixed pairs and hybrids (Mazgajski & Rejt 2006; Ciach & Fröhlich 2013), as the methodology used (typically fast counts based on only acoustic detection) does not allow for the discovery of hybrids (Fröhlich et al. 2022).Consequently, the results of most studies are at least uncertain if not simply erroneous, as the data omit hybrid individuals and wrongly assign mixed pairs to one of the species (usually GW as it is generally more abundant). Error assignment of individuals to a “pure” species cannot be neglected as such birds have some intermediate biological, ecological and ethological properties. This is particularly visible regarding habitat preferences, which only partially overlap between GW and SW (Figarski & Kajtoch 2018). In addition, reproduction could be different as GW usually starts to breed earlier than SW (Wesołowski & Tomiałojć 1986; Michalczuk & Michalczuk 2016), which is less adapted to the harsh winters of Central and East Europe. These problems affect also studies on GW breeding in urban forests surrounded by urban woodlands occupied by SW, thus interactions and hybridization of these species are likely in the contact zone of these environments.

### 4.6. Recording and monitoring of woodpeckers

Difficulties in detecting hybrid woodpeckers can result in adverse consequences for the recording and monitoring of woodpeckers for demographic and/or conservation purposes. Detection based only on responses to the playback of calls and drumming can be misleading as the sounds made by hybrids have not been adequately studied (Fig. 7). Winkler and Short (1978) suggested that some intermediate calls of hybrids exist, but the personal observations of the authors of this paper suggest that the calls and drumming of hybrids can be similar to one of the parental species (perhaps depending on the share of genomes or on the imprinting of calls from one of the parents). The question also arises whether hybrid individuals can change their calls to resemble either GW or SW depending upon which species it is responding to. It is unclear if calls and drums are merely inherited or are learned during the immature phase of growth. The interspecific origin of hybrids makes the development of calls and drumming an interesting topic of study as it is not fully understood. Therefore, only direct observations (combining calls, drums and visual detection) allow for the determination of such hybrids. Furthermore, often only photographs can result in a reliable assignment of an individual as a hybrid. Therefore, monitoring conducted without these guidelines should not be treated as dependable. Overall, the methods used in monitoring common breeding birds in Europe are unfortunately unsuitable for the detection of hybrids. These methods (without using playback) are even inappropriate for the detection of SW as this species is often hard to detect if playback is not used (Michalczuk 2006). According to our knowledge, there are no specific programmes that enable the monitoring of SW populations in the European Union and the estimated population trends for this species in most countries are simply based on the knowledge of local birders. Notwithstanding the expertise of such observers, this situation is clearly not ideal. Certainly, this is the case in Poland, where a population of SW is assumed to be stable over the last 20-30 years, while repeated counts with more appropriate methods revealed a 40% decrease in number in rural and urban populations (Michałczuk & Michałczuk; Kajtoch & Kusal 2023). Currently, almost nothing is known about changes in interspecific reproduction and hybrid occurrence. Therefore, there is a need to implement a suitable monitoring programme for SW and hybrids in the European Union (and Europe as a whole), at least for populations on the edge of the SW range.

**Fig. 7.**
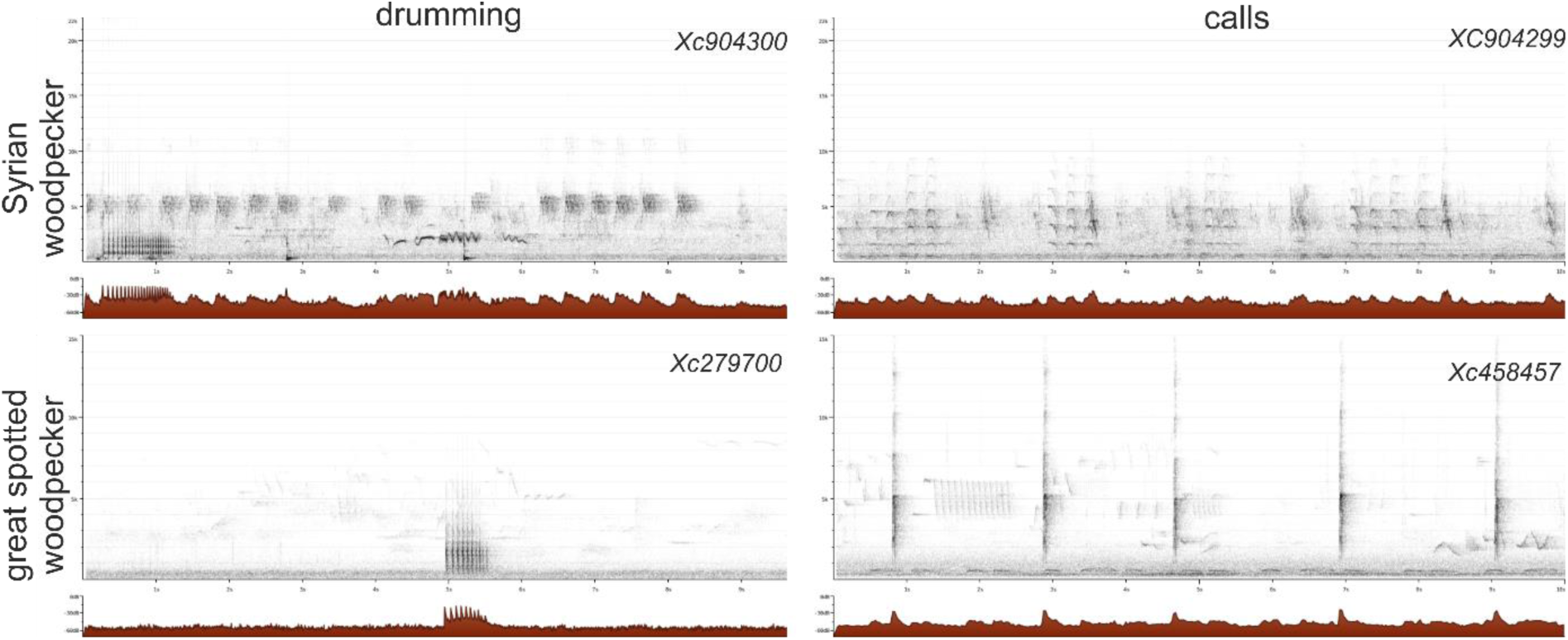
Drumming and calls of Syrian and great spotted woodpeckers. Sonograms from xeno- canto: David Darrell-Lambert, XC458457. Accessible at www.xeno-canto.org/458457, David Darrell-Lambert, XC904299. Accessible at www.xeno-canto.org/904299, David Darrell-Lambert, XC279700. Accessible at www.xeno-canto.org/279700, David Darrell-Lambert, XC904300. Accessible at www.xeno-canto.org/904300)

### 4.7. Perspectives and recommendations

The co-occurrence of SW and GW opens up many opportunities for research that can extend our knowledge of species interactions and hybridisation. Many questions concerning sympatric populations of these woodpeckers are still unanswered. Recently, ecology and ethology issues were examined (Gorman 2015, Michałczuk & Michałczuk 2016, 2017), but still, most of the data refers to “pure” birds, and there is a need to extend this to mixed pairs and hybrids in order to resolve the hypothesis that such birds have intermediate preferences. Even the plumage of hybrids requires more detailed study as it is unknown whether there is one “typical” appearance of such birds or if characteristics are different depending on the share of parental taxa genomes. There is at present scant knowledge on the reproductive biology of mixed GW and SW pairs and pairs composed of existing hybrids, although it has been proven that they are fertile (Kajtoch & Kusal 2023). It is unknown whether hybrids and backcrosses are equally as viable in terms of similar lifespans, adaptations to the environment and breeding success, as “pure” birds. It is possible that hybrids will in further generations have lower fitness levels. Another key question is whether hybrid populations could ultimately lead to these two woodpeckers losing their “species specific” status. This is unlikely over the whole ranges of GW and SW, which are mostly allopatric, but could result in the formation of local “hybrid swarms” (Dufresnes et al. 2018). This is one of topics currently being examined under the project led by the Institute of Systematics and Evolution of Animals of the Polish Academy of Sciences in cooperation with several scientific institutes and ornithological organisations in Europe (for further details see: https://sites.google.com/view/hybridization-woodpeckers/). Such local hybrid populations could have consequences for the conservation of SW, especially if the decline in some areas extends to other regions (Winkler et al. 2014). Finally, the question remains whether hybrid populations should be protected as “pure” ones are. Unfortunately, current European Union nature and conservation legislation neglects this natural phenomenon. For the understanding and proper management of SW and also hybrid SW x GW populations, we recommend that a monitoring programme, using appropriate methodology, is implemented.

## Acknowledgements

This review was prepared as a part of the grant funded by National Science Center, Poland (project number UMO-2022/47/O/NZ9/02044, granted to Ł. Kajtoch).

Presented manuscript is fully based on already published data, therefore no permissions (for study protected species or conducting research in protected areas from any country) were required.

## Statements and Declarations

### 1. Compering Interests

The authors have no competing interests to declare that are relevant to the content of this article.

### 2. Ethical

No approval of research ethics committees was required to accomplish the goals of this study because there was no experiments conducted in this review study.

### 3. Author contributions

All authors contributed to the study conception and design. The original idea belongs to Łukasz Kajtoch, data analysis and literature search was performed by Antonii Bakai. Gerard Gorman provided main critical review of the work. The first draft of the manuscript was written by Antonii Bakai and all authors commented on previous versions of the manuscript. Gerard Gorman shape original text in to grammatical accurate English. All authors read and approved the final manuscript.

---

**Table S1.**
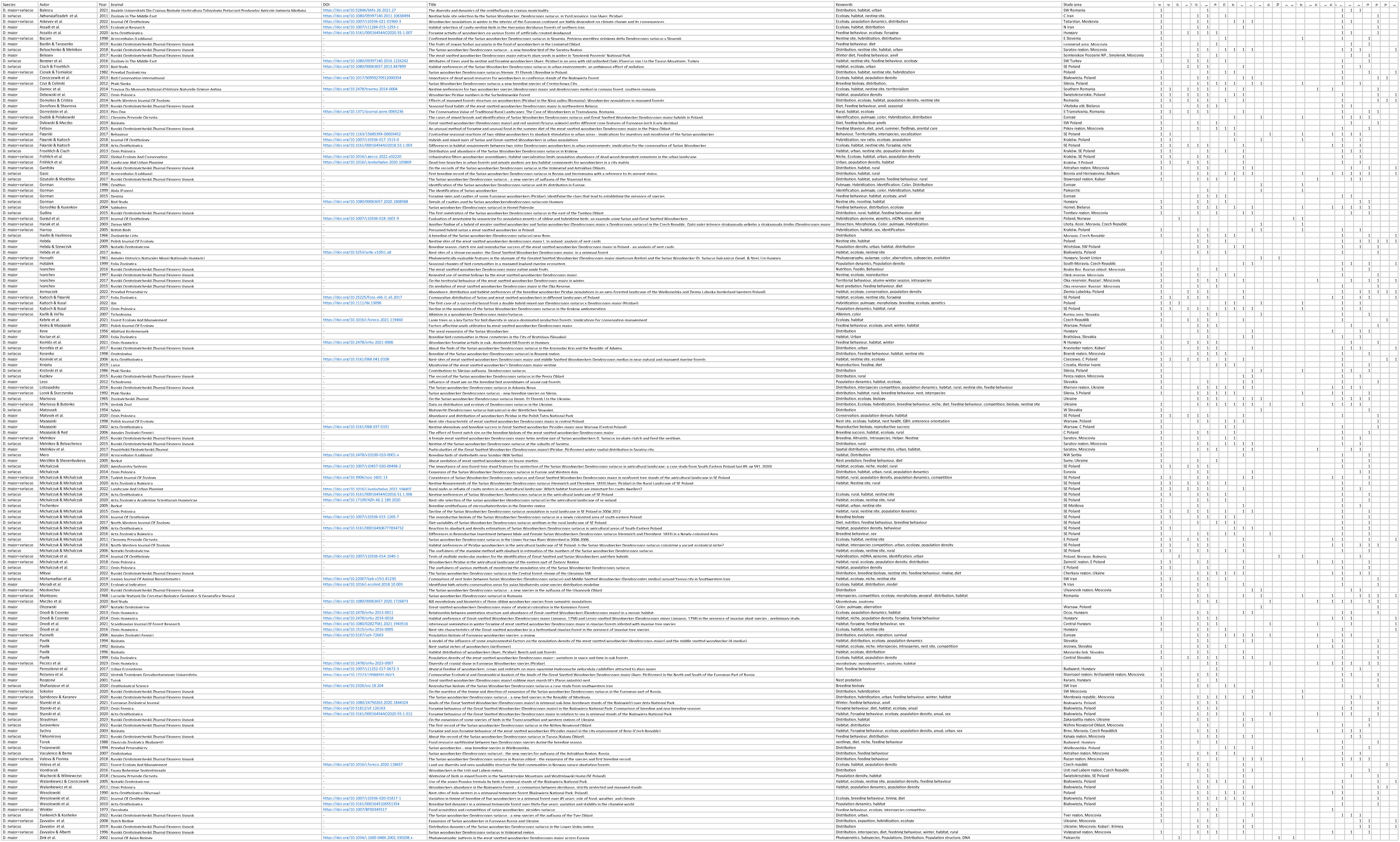
Summary of literature search in Web of Science and Scopus databases for articles on Syrian and great spotted woodpeckers in their sympatric range.

